# MTH1 inhibitor TH588 induces mitosis-dependent accumulation of genomic 8-oxodG and disturbs mitotic progression

**DOI:** 10.1101/573931

**Authors:** Sean G Rudd, Helge Gad, Nuno Amaral, Anna Hagenkort, Petra Groth, Cecilia E Ström, Oliver Mortusewicz, Ulrika Warpman Berglund, Thomas Helleday

**Author notes:** Corresponding author. Mailing address: Department of Oncology and Metabolism, Medical School, S10 2RX Sheffield, UK.

## Abstract

Reactive oxygen species (ROS) oxidise nucleotide triphosphate pools (e.g., 8-oxodGTP), which may kill cells if incorporated into DNA. Whether cancers avoid poisoning from oxidised nucleotides by preventing incorporation via the oxidised purine diphosphatase MTH1 remains under debate. Also, little is known about DNA polymerases incorporating oxidised nucleotides in cells or how oxidised nucleotides in DNA become toxic. We show replacement of one of the main DNA replicases in human cells, DNA polymerase delta (Pol δ), to an error-prone variant allows increased 8-oxodG accumulation into DNA following treatment with the MTH1 inhibitor (MTH1i) TH588. The resulting elevated genomic 8-oxodG correlates with increased cytotoxicity of TH588. Interestingly, no substantial perturbation of replication fork progression is observed, but rather mitotic progression is impaired and mitotic DNA synthesis triggered. Reducing mitotic arrest by reversin treatment prevents accumulation of genomic 8-oxodG and reduces cytotoxicity of TH588, in line with the notion that mitotic arrest is required for ROS build-up and oxidation of the nucleotide pool. Furthermore, we demonstrate delayed mitosis and increased mitotic cell death following TH588 treatment in cells expressing the error-prone Pol δ variant, which is not observed following treatments with anti-mitotic agents, thus linking incorporation of oxidised nucleotides and disturbed mitotic progression.

## INTRODUCTION

Oxygen metabolism is critical for many cellular processes, and it is established that reactive oxygen species (ROS) can be both beneficial and deleterious to cells. Loss of balanced redox homeostasis and/or ROS may have a role in the etiology of many diseases, including cancer, cardiovascular disease, hypertension, inflammatory diseases (e.g., atherosclerosis, rheumatoid arthritis), ischemia-reperfusion injury, diabetes mellitus, neurodegenerative diseases, and ageing (1,2). ROS causes oxidative DNA damage, either directly in the DNA, which then can be repaired by the 8-oxoguanine glycosylase OGG1, or by oxidising the nucleobases in the free deoxynucleoside triphosphate (dNTP) pool. The human MutT homologue 1 (MTH1) protein, an oxidised purine diphosphatase, is important to prevent oxidised dNTPs, such as 8-oxodGTP and 2-OHdATP, from being incorporated into DNA (3,4).

Previously, we and others showed that the normally non-essential MTH1 enzyme is required for survival of cancer cells (5–8), while others have ruled out MTH1 as an anti-cancer target altogether (9–11). We showed that the wild-type MTH1 or bacterial MutT protein can rescue 8-oxodGTP incorporation and toxicity generated following MTH1 siRNA depletion and treatment with the MTH1 inhibitor (MTH1i) TH588 (6,12), consistent with incorporation of 8-oxodGTP into DNA having toxic effects. MTH1i TH588 and the best-in-class analogue, the clinical candidate TH1579 (karonudib), possess dual pharmacology, inhibiting microtubule polymerisation directly in addition to inhibiting MTH1 (10) and the MTH1 protein itself is required for mitosis and binds tubulin directly (Gad et al., submitted), which altogether are complicating the understanding of what effects are generated through mitosis or through incorporation of 8-oxodGTP.

Pol δ is a main DNA polymerase involved in lagging strand synthesis as well as being the main DNA polymerase for repair or break-induced replication (13). Emerging data also suggests Pol δ is the main DNA polymerase for leading strand synthesis (14). Unlike trans-lesion synthesis polymerases, Pol δ has less tolerance to replicate across lesions and only accepts very slight base modifications to be able to polymerise DNA with modified nucleotide substrates. Hence, 8-oxodGTP is likely a poor substrate for Pol δ in cells. To investigate the cellular fate of 8-oxodGTP, we here use a recently reported protein-replacement system allowing the replacement of endogenous Pol δ with an error-prone variant (15). Here, we report increased incorporation of 8-oxodGTP when cells express the error prone variant are exposed to MTH1i TH588 enabling us to use this model to increase our understanding of the role of genomic 8-oxodG lesions. These experiments are highly relevant for cancer treatment as MTH1i TH1579 (karonudib) is currently evaluated in clinical trials for anti-cancer efficacy.

## MATERIALS AND METHODS

### Cell culture

All cell lines were cultured at 37°C in 5% CO2. U2OS cells (ATCC) were cultured in DMEM media supplemented with 10% FBS and Penicillin Streptomycin (100 U/ml), A2780 Pol δ replacement cells (15) were cultured in DMEM with 5% low TET FBS and 100 μg/mL G418, 10 μg/mL Blasticidin and 1 μg/mL Puromycin. All cell culture reagents were from Gibco/Thermo Fisher.

### Compounds

The following chemical inhibitors were used: CENP-E inhibitor (GSK923295, SelleckChem), Vincristine sulfate (Sigma Aldrich), Paclitaxel (Sigma Aldrich), Reversine (Axon MedChem), RO3306 (SelleckChem), Aphidicolin (Sigma Aldrich). TH588(6) was synthesized in house and prepared as previously reported.

### Antibodies

The following antibodies were used: mouse anti beta-Actin (ab6276, Abcam), mouse anti-ATM pS1981 (sc47739, Santa Cruz Biotech), rabbit anti-ATR phospho S428 (ab178407, Abcam), rabbit anti-Cdk2 phospho-T14/T15 (Santa Cruz, sc28435-R), rabbit anti-Chk1 phospho S345 (#2341, Cell Signaling), mouse anti-Chk1 (#2360, Cell Signaling), rabbit anti-GAPDH (sc-25778, Santa Cruz), mouse anti-H2A.X phospho S139 (05-636, Millipore), mouse anti-Histone H3 phospho-S10 (H3-pS10; ab14955, Abcam), rabbit anti-Histone H3 (#4318, Cell Signaling), rabbit anti-MTH1 (NB100-109, Novus Biologicals), rabbit anti-p21 (H164, Santa Cruz Biotech), rabbit anti-cleaved PARP (#9541, Cell Signaling), mouse anti-PLK1 (Millipore, #5844), mouse anti alpha-Tubulin, mouse anti-DNA Pol δ catalytic subunit (p125) (ab10362, Abcam).

### Generation of H2B-GFP cell lines

The H2B-GFP vector was constructed by amplifying H2B from the H2B-RFP pENTR1A vector (Addgene # 22525) by PCR and subcloning the product into the pENTR1A-GFP-N2 vector (Addgene # 9364) at the HindIII and BamH1 restriction sites. The H2B-pENTR1A-GFP-N2 vector was verified by sequencing and transferred into the pLenti-CMV-Blast and pLenti-CMV-Hygro vectors using LR clonase (Invitrogen). H2B-RFP in pENTR1A (w507-1) and pENTR1A-GFP-N2 (FR1) were gifts from Eric Campeau and Paul Kaufman (Addgene plasmids # 22525 and # 19364). H2B-GFP plasmids were transduced using lentivirus infections.

### Western blot

Cells were washed with 1x PBS and scraped in lysis buffer (50 mM Tris-HCl pH 8, 150 mM NaCl, 1 mM EDTA, 1% Triton X-100, 0.1% SDS, 1x protease inhibitor cocktail (Thermo), and 1x Halts phosphatase inhibitor cocktail (Thermo)). Samples were incubated on ice for 30 min with occasional vortexing before centrifugation to pellet insoluble material. Protein concentration was measured using BCA Assay (ThermoFisher) before preparation in Laemmli Sample Buffer (Bio-Rad) containing 355 mM final concentration 2-mercaptoethanol, and samples were denatured at 95°C for 5 min. Samples were run on 4-12% SDS-PAGE gels (Mini-Protean TGX Precast gel, Bio-Rad) and subsequently transferred to nitrocellulose membranes using the Trans-Blot Turbo instrument (Bio-Rad). Membranes were stained with Ponceau S and blocked in Odyssey TBS Blocking Buffer (Li-Cor) for 1 h. Antibodies were diluted in Odyssey TBS Blocking Buffer and incubated overnight at 4°C. Membranes were washed with TBS supplemented with Tween-20 (0.1%) before incubation with secondary antibodies (IRDye 800CW and IRDye 680LT) diluted in TBS-Tween before further washes and imaging using the LI-COR Odyssey Fc Imaging system.

### Cell viability assay

Cell viability was assessed using the resazurin assay. In the case of Pol δ replacement cells, prior to the cell viability assay, cells were seeded in 6-well plates and cultured in medium supplemented with 10 ng/ml doxycycline for 3 days before trypsinization and re-seeding; doxycycline supplementation was maintained for the remainder of the assay. For the cell viability assay, typically 1500 cells per well were seeded in 90 μl media in 96-well plates, and the following day, the drug to be tested was serially diluted in media and 10 μl 10x drug dilution added to the desired well. Following 3 days treatment, viable cells were measured using resazurin (Sigma Aldrich; cat no. R7017), as described previously (6) Measurements for each well were normalized to control wells (cells with DMSO, 100% viable control; medium with DMSO, 0% viable control) and analyzed using four-parameter logistic model in Prism 8 (GraphPad Software).

### DNA fiber technique

Cells were seeded in 6-well plates, typically 200 000 – 300 000 per well, to reach 70% confluency on day of experiment. For the time-course analysis in U2OS cells, 5 μM TH588 was added 8, 4, and 1 hour before labeling or directly with labeling, and an 8 hour DMSO control was also included. For experiment with the Pol δ replacement cells, cells cultured in 10 ng/ml doxycycline for 4 days were treated with 1 μM TH588 or mock treated with DMSO for the indicated times. Treatment was followed by pulse-labelling for 20 minutes with 25 μM 5-Chloro-2′-deoxyuridine (CldU, Sigma) and then 20 minutes with 250 μM 5-Iodo-2′-deoxyuridine (IdU, Sigma). The inhibitor and mock treatment was present during labelling. Cells were harvested and resuspended in ice-cold PBS. Lysing of cells were performed by incubating a drop of cell suspension together with spreading buffer (200 mM Tris–HCl, pH 7.4, 50 mM EDTA and 0.5% SDS) on SuperFrost objective glass (Thermo Scientific). DNA fibers were spread by tilting the glass slides allowing the cell lysate to slowly run down the slide. For immunodetection of CldU and IdU, acid treated fibers (2.5 M HCl for 1 hour) were stained with monoclonal rat anti-BrdU (1:500) (Clone BU1/75 (ICR1) (AbD Serotec) and monoclonal mouse anti-BrdU (1:500) (Clone B44, 347580 (BD Biosciences)) respectively for 1 hour in 37°C. This was followed by incubation with Alexa555-conjugated anti-rat and Alexa488-conjugated anti-mouse (Molecular Probes). DNA fibers were studied in a LSM780 confocal microscope using a 64x oil objective and lengths of CldU and IdU tracks were measured using the ImageJ software (http://fiji.sc/Fiji), 50-100 replication tracks were scored for each condition.

### Mitotic DNA replication

U2OS cells were seeded on coverslips one day before. The next day the cells were incubated for 16h with the CDK inhibitor RO3306 (9μM) alone or in combination with either 0.4μM Aphidicolin (APH), 5μM TH588, or 500nM TH1579. Afterwards the cells were washed with pre-warmed PBS 4-5 times and within 5 minutes to release the cells into mitosis and incubated with fresh media containing 10μM EdU for 30min at 37ºC. Fixation and permeabilisation was done simultaneously with PTEMF solution for 20min at room temperature. Blocking was done in 3% BSA in PBST 0.5% for 1h at room temperature followed by EdU detection with the Click-iT EdU reaction kit (Thermo Scientific), following the manufacturer’s instructions. The nucleus was visualized with DAPI. For labelling of mitotic cells anti-H3pS10 primary antibody was used.

### High content microscopy

Pol δ replacement cells were first cultured with media supplemented with 10 ng/ml doxycycline for 3 days before seeding at 10 000 cells per well in black, clear-bottomed 96-well plates (BD Falcon). The following day, if labelling S-phase cells, cells were pulse-labelled with 10 μM EdU for 20 minutes prior to fixation in 4% paraformaldehyde in PBS (Santa Cruz) containing 0.5% Triton X-100 for 15 min. Wells were washed with PBS before permeabilisation in 0.3% Triton X-100. For detection of EdU labelled cells, a click reaction mix was assembled in PBS containing 2.5 mM copper (II) sulphate, 10 mM ascorbic acid and 2 μM ATTO-488 azide fluorophore (ATTO-TEC), which was added to wells and incubated in the dark for 30 minutes. Wells were washed in PBS, cell nuclei stained with 1 μg/ml DAPI for 10 min, and further washed in PBS prior to imaging. Plates were imaged on an ImageXpress high-throughput microscope (Molecular Devices) with a 20x objective. Images were analyzed (nuclei counting, DAPI and EdU nuclear intensity measurements) with CellProfiler (Broad Institute) and data handled in Excel and plotted in Prism6.

### Modified comet assay

Cells were seed in 6-well plates at a density of 150-200 000 cells per well and either on the same day (suspension cells) or the day after (adherent cells) treated for 24h with compound or DMSO control. Cells were harvested by trypsinization, washed once in PBS and resuspended in 300μl PBS. The suspension (100 μl) was mixed with 500 μl 1.2% low melting point agarose at 37°C and the mixture were added to agarose coated slides and a coverslip was added on top. The slides were solidified on ice and lysed O/N at 4°C in Lysis buffer (2.5 M NaCl, 100 mM EDTA, 10 mM Tris, 10% DMSO, 1% Triton X100). Samples were washed 3 times in Enzyme buffer (40 mM HEPES, 0.1 M KCl, 0.5 mM EDTA, 0.2 mg/ml BSA, pH 8.0) and treated with hOGG1 enzyme (2 μg/ml) or buffer alone for 45 min at 37°C. hOGG1 was expressed in *E. coli* and purified as previously described (Gad et al 2014). Slides were transferred to Alkaline Electrophoresis buffer (300 mM NaOH, 10 mM EDTA) for 30 min and electrophoresis was performed at 25V, 300 mA for 30 min at 4°C. Samples were washed in 400 mM Tris pH 7.5 for 45 min. DNA was stained with SybrGOLD and comets were quantified with Comet Assay IV software in live video mode.

### Time-lapse microscopy

Cells were seeded in 96-well plates (BD Falcon plate 353376; 5000 cells/well). The day after, cells were treated and the time-lapse was initiated after 30 min and images of GFP and brightfield channels were acquired every 10 min for 24 h in a Perkin-Elmer ImageXpress instrument. Cells were kept at 37°C in 5% CO_2_ atmosphere during the entire time-lapse experiment. Movie files were assembled in MetaXpress and ImageJ softwares. For each condition, images were acquired from two different wells. Individual cells were followed manually and scored for defects in mitosis including mitotic slippage/polynucleation (MS/PN), micronuclei formation (G1/MN), mitotic slippage (MS) or cell death during mitosis (DiM). The time in mitosis was defined as the time from the cell rounding up and condensing the chromatin to the end of cytokinesis.

### Statistical analysis

All statistical analysis was performed using Prism 8 (GraphPad software).

## RESULTS

### An experimental system to study 8-oxodGTP incorporation into DNA

Our original model on the role of MTH1 in protecting cancer cells centred upon the mis-incorporation of oxidised nucleotides, such as 8-oxodGTP, into DNA as the toxic lesion (6). More recently, others and ourselves have demonstrated direct action of TH588 on tubulin polymerisation and that MTH1 itself has a novel role in mitosis, rendering a more complex mechanism of action of TH588 and MTH1 as an anti-cancer target (Gad et al., submitted). With this background it is important to understand if 8-oxodG in DNA is a potentially lethal lesion and how this is incorporated in cells. To re-examine this model we employed a recently reported protein-replacement system that allows the doxycycline-inducible replacement of endogenous Pol δ, one of the main DNA replicases in cells, with either a wild type (WT), error-prone (EP, L606G), proofreading-deficient (PD, D402A), or EP and PD double mutant (DM) variant in the ovarian cancer cell line A2780 (Figure 1a). We were particularly interested in the EP variant as we reasoned this would increase mis-incorporation of oxidised nucleotides into DNA and thus provide a controlled experimental system to study the consequences of TH588-treatment. Importantly, within the timeline of the experiments performed, replacement of endogenous Pol δ with the Pol δ variants did not significantly reduce cell viability (Figure 1b) and had a minimal effect on the proportion of S-phase cells (Figure 1c), with the exception of the DM variant, which had a small but significant increase in the proportion of S-phase cells, as previously reported (15). To determine whether replacement of Pol δ with the EP variant and subsequent TH588 treatment leads to increased genomic 8-oxodG, we employed the modified comet assay in which cells embedded in agarose are exposed to recombinant OGG1 prior to electrophoresis under alkaline conditions. The presence of 8-oxodG in DNA would lead to increased OGG1-induced comet tails in the assay. Consistent with our previous reports (6,16), TH588 treatment, irrespective of Pol δ status, resulted in increased 8-oxodG in DNA as compared to DMSO treated controls (Figure 1d, e). The highest accumulation of genomic 8-oxodG was observed in the TH588-treated cells expressing the Pol δ EP variant (Figure 1d, e), indicating that this reduced-fidelity variant of this replicase mis-incorporates 8-oxodG into DNA. Thus, we have an experimental system with which to study the cellular consequences of increased mis-incorporation of 8-oxodGTP into DNA.

**Figure 1.**
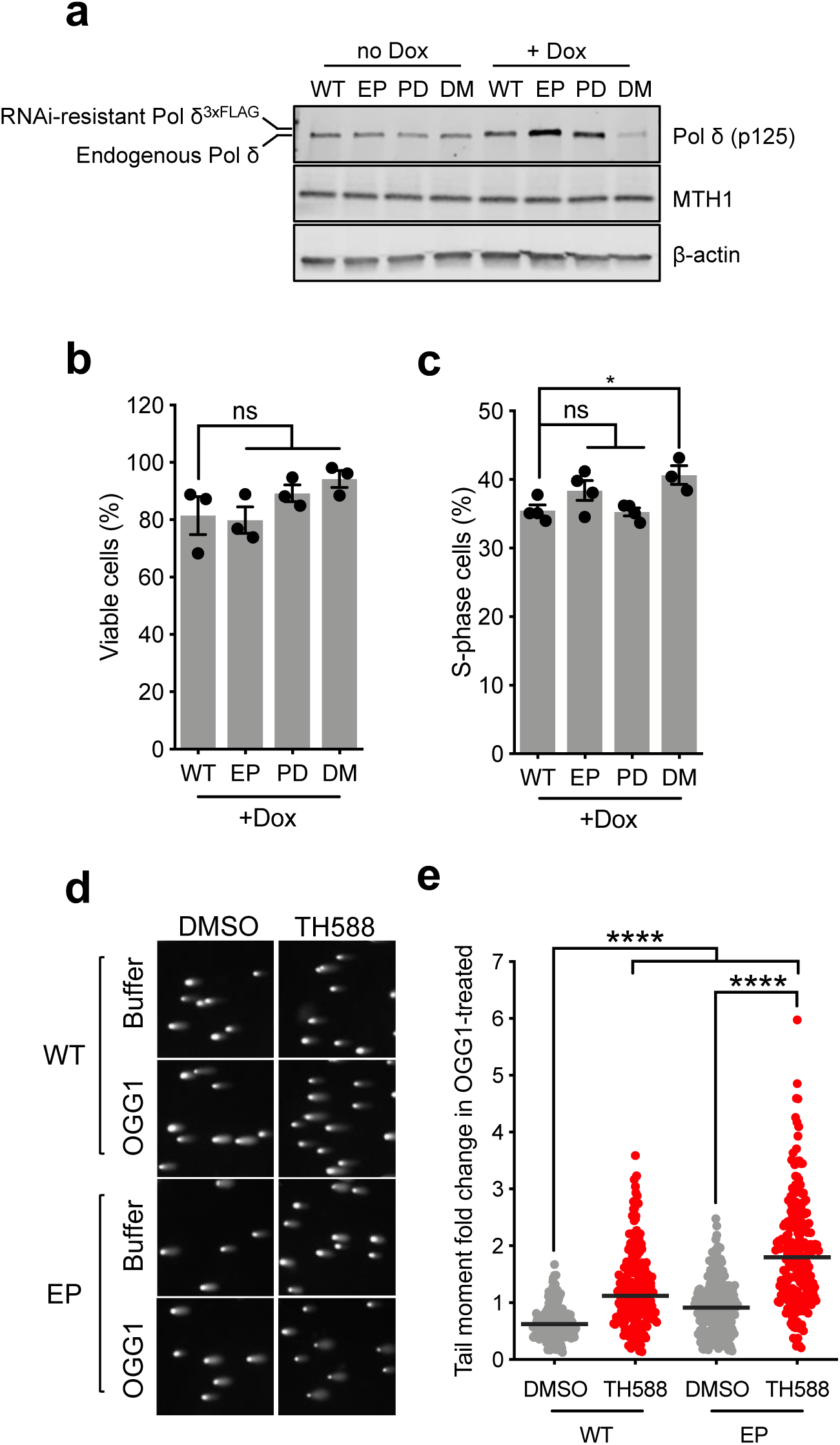
An experimental system to study 8-oxodGTP incorporation into DNA. **a)** Pol δ replacement cells (WT, wild type; EP, error-prone; PD, proofreading deficient; DM, EP and PD double mutant) were cultured in the presence or absence 10 ng/ml doxycycline for 4 days and lysates prepared for western blot analysis with the indicated antibodies. **b)** Pol δ replacement cells were cultured in 10 ng/ml doxycycline for 7 days and viable cells measured using the resazurin assay. Values were normalized to Pol δ replacement cells cultured in the absence of doxycycline. Averages of 3 experiments shown each performed in sextuplet, error bars indicate SD. **c)** Pol δ replacement cells were cultured in 10 ng/ml doxycycline for 4 days were labeled with 10 μM EdU for 20 minutes prior to fixation. S-phase cells were defined at >9 EdU foci per nucleus, at least 2000 cells were analysed per well with experiments performed at least with duplicate wells. Average of at least 3 independent experiments are plotted, error bars indicate SD. **d, e)** Pol δ replacement cells (WT, wild type; EP, error prone) were cultured in 10 ng/ml doxycycline for 4 days before treatment with 1 μM TH588 for 24 hours. Genomic levels of 8-oxo-dG were analysed using the modified comet assay, representative images are shown in (d). Fold difference in tail moment comparing OGG1-treated to buffer only control shown in (e). At least 200 cells were analysed in 2 independent experiments, horizontal bar indicates the median, statistical comparison performed using Mann-Whitney tests (****, p<0.0001).

### Increased genomic 8-oxodG in TH588-treated Pol δ EP cells correlates with increased toxicity

The cytotoxic effects of TH588 have been proposed to be independent of targeting MTH1 and independent of 8-oxodGTP (9,10,17), while we have shown TH588 cytotoxicity is related to ROS and partly rescued by expression of the bacterial 8-oxodGTPase MutT (6,12,18). Here, we wanted to use this experimental system to determine the contribution of 8-oxodGTP incorporation to the cytotoxicity of TH588. In accordance with the original proposed mechanism of action of MTH1 inhibition, replacement of endogenous Pol δ with the EP but not WT variant increased TH588-induced cytotoxicity (Figure 2a). Elevated sensitivity was also observed in the cells expressing the Pol δ DM but not PD variant (Figure 2a), indicating the increased sensitivity was due to the point mutation reducing Pol δ fidelity, i.e. the EP variant. In line with this, elevated DNA damage (γH2Ax) and apoptotic (cleaved-PARP) signalling was observed in the EP and DM variant expressing cells compared to WT control (Figure 2b). Taken together, this data indicates that incorporation of TH588-induced 8-oxodGTP contributes to cell killing.

**Figure 2.**
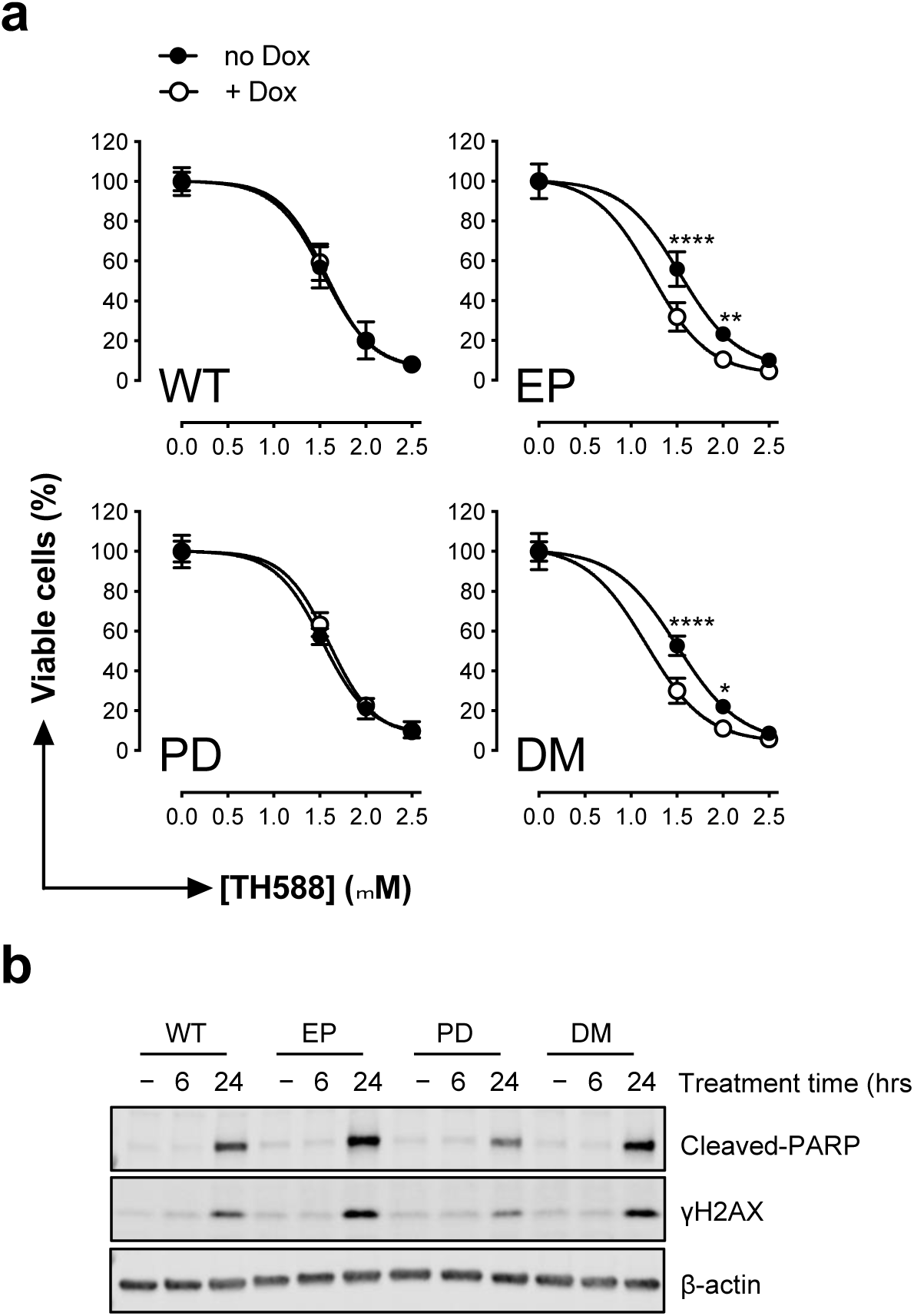
An error-prone variant of Pol δ sensitizes cells to TH588. **a)** Pol δ replacement cells (WT, wild type; EP, error-prone; PD, proofreading deficient; DM, EP and PD double mutant) were cultured in the presence or absence 10 ng/ml doxycycline for 3 days prior to re-seeding, and 24 hours later, treatment with a dose-response of TH588 in media with or without doxycycline. Viable cells were measured following a 3 day treatment. A representative of 4 independent experiments is shown, performed in sextuplet, with error bars indicating SD. Statistical comparisons performed with a two-way ANOVA (*, p≤0.05; **, p≤0.01; ****, p<0.0001). **b)** Pol δ replacement cells were cultured with doxycycline for 4 days and then treated with 1 μM TH588 for the indicated times before collection for western blot analysis with the indicated antibodies. Representative blot of 2 independent experiments shown.

### 8-oxodGTP incorporation does not greatly perturb S-phase DNA synthesis

It is easy to envision that mis-incorporation of 8-oxodGTP would generate replication stress, as MTH1 overexpression reduces Ras-induced DNA lesions and counteracts oncogenic stress (8,19), and replication stress is associated with oxidative stress in cancer (20) and replication difficulties (21,22). Therefore, we wanted to investigate if MTH1 inhibition and resulting incorporation of oxidised nucleotides would result in replication stress. To test this we used the well-established DNA fibre assay in cells expressing either the Pol δ WT or EP variant treated with TH588 for 0-6 hours prior to the assay. Two hours of TH588 treatment resulted in a small, but significant, decrease in replication fork speeds regardless of whether cells were expressing the WT or EP variant of Pol δ, and this decreased fork speed recovered to DMSO-treated control levels following 6 hours TH588 treatment (Figure 3a, b). Notably, expression of the Pol δ EP variant did not significantly alter this TH588-induced replication defect compared to WT controls (Figure 3a, b). In a parallel set of experiments, the replication stress response was also monitored in osteosarcoma U2OS cells treated with TH588, which we have previously reported are sensitive to TH588 treatment and accumulate genomic 8-oxodG (6). Similarly, an initial small, but significant, decrease in replication fork speeds were observed following TH588 treatment, but fork speeds quickly recovered to that of the control following longer treatment times (**Supplemental Figure 1a, b**). Consistent with no great perturbation of ongoing DNA synthesis, no activation of the intra-S-phase checkpoint kinase Chk1 was observed following TH588 treatment, despite the induction of DNA damage signalling at later timepoints (**Supplemental Figure 1c)**.

**Figure 3.**
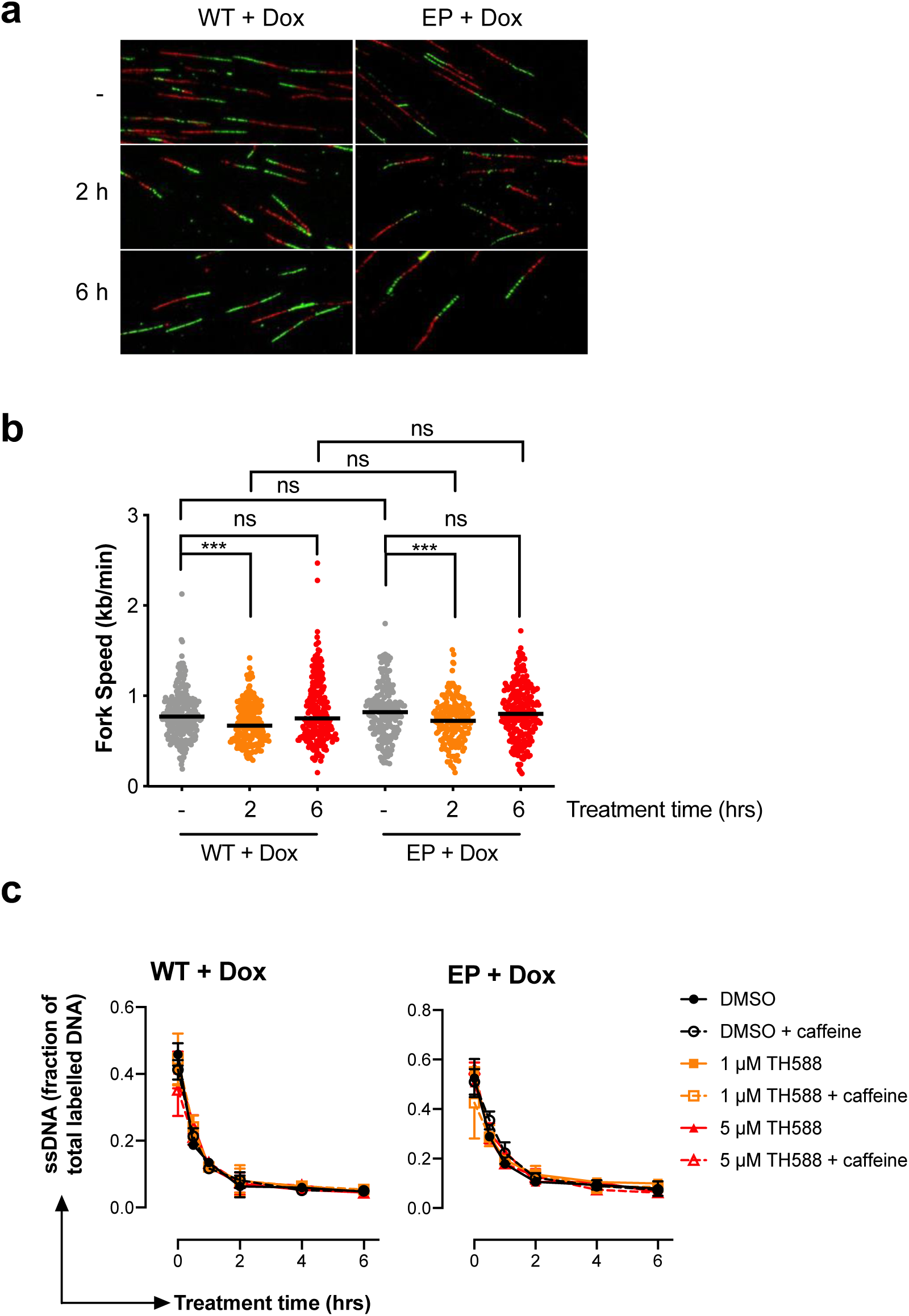
Incorporation of TH588-induced 8-oxodGTP does not cause sustained replication stress in S-phase. **a, b)**. Pol δ replacement cells were cultured with doxycycline for 4 days before treatment with 1 μM TH588 for the indicated time and sequential pulse labeling with CldU and IdU prior to preparation and analysis of spread DNA fibres. Representative DNA fibre images are shown in (a). Measured fork speeds from two independent experiments are shown in (b), with at least 200 fibers measured in total. Horizontal pars indicate the median, statistical testing was performed using Mann-Whitney tests (ns, not significant; **, p≤0.01) **C)** Alkaline DNA unwinding assay with Pol δ replacement WT or EP cells treated with 10 ng/ml doxycycline for 4 days prior to the indicated treatments. Average from 2 independent experiments shown, mean values plotted with error bars indicating SD. Caffeine is used as the post-replication repair of gaps is caffeine dependent, hence, the detection of gaps can be increased with caffeine.

Upon DNA damage, replication may proceed downstream of a stalled fork by origin-independent re-priming, e.g., by PrimPol (23,24), leaving a gap on the nascent DNA strand which cannot be detected by the DNA fibre assay (25). To test if gaps are generated along the replicating nascent DNA strand, we made use of an in house assay which detects such gaps (26). Briefly, we ^3^H-thymidine pulse-labelled ongoing forks, and then let these progress from the labelled area for different times. By addition of alkaline solution, DNA unwinding takes place from the single-stranded ends of the fork or gaps, if present. Hence, as the fork moves forward or the gaps are filled, the labelling is removed from the fraction of DNA that becomes single-stranded. Using this assay we demonstrate that no gaps are generated behind the fork during replication following TH588 treatment, regardless of Pol δ status (Figure 3c). Taken together, these data indicate that despite incorporation of oxidised nucleotides, and the ultimate cytotoxicity, no substantial, sustained perturbation of ongoing DNA synthesis in S-phase cells is observed following TH588 treatment.

### Treatment with TH588 triggers mitotic DNA replication

Emerging data shows TH588 and MTH1 siRNA arrest cells in mitosis through interference of tubulin dynamics, and in the case of TH588, primarily due to direct inhibition of tubulin polymerisation (10) (Gad et al., submitted). The explanation for MTH1i or MTH1 siRNA to fail to be toxic in cancer cell lines could be explained by lack of ROS, which is supported by non-toxic MTH1i becoming toxic in the presence of ROS and also that MTH1 knockout zebrafish embryos are sensitive to microinjection of 8-oxodGTP (Gad et al, submitted). It has been reported that ROS and oxidative stress are elevated during mitosis and that mitotic arrest itself generates ROS (27), with one report linking this to mitophagy (28), and hence MTH1 may have a role in hydrolysing oxidised purine nucleotides during mitosis. Normally, DNA replication is completed during the S-phase of the cell cycle. However, under conditions of replication stress, such as that generated experimentally by low-dose aphidicolin treatment, it has been demonstrated that under-replicated regions are repaired by Pol δ–mediated mitotic DNA replication at common fragile sites (29). Since replication stress is often found in cancer (21), this may explain how 8-oxodGTP could be specifically incorporated in cancer cells following MTH1 siRNA or TH588 treatment, as previously reported. To test if TH588 would induce mitotic DNA replication, we synchronized U2OS cells in G2 using the Cdk1 inhibitor RO-3306 together with 0.4 μM aphidicolin as a positive control or TH588. Cells were released into mitosis for 30 min by washout of the compounds and EdU was added to the media to label newly synthesized DNA. As expected, aphidicolin induced EdU foci in mitotic cells and so did TH588 (Figure 4a, b), suggesting that mitotic replication is stimulated with this compound.

**Figure 4.**
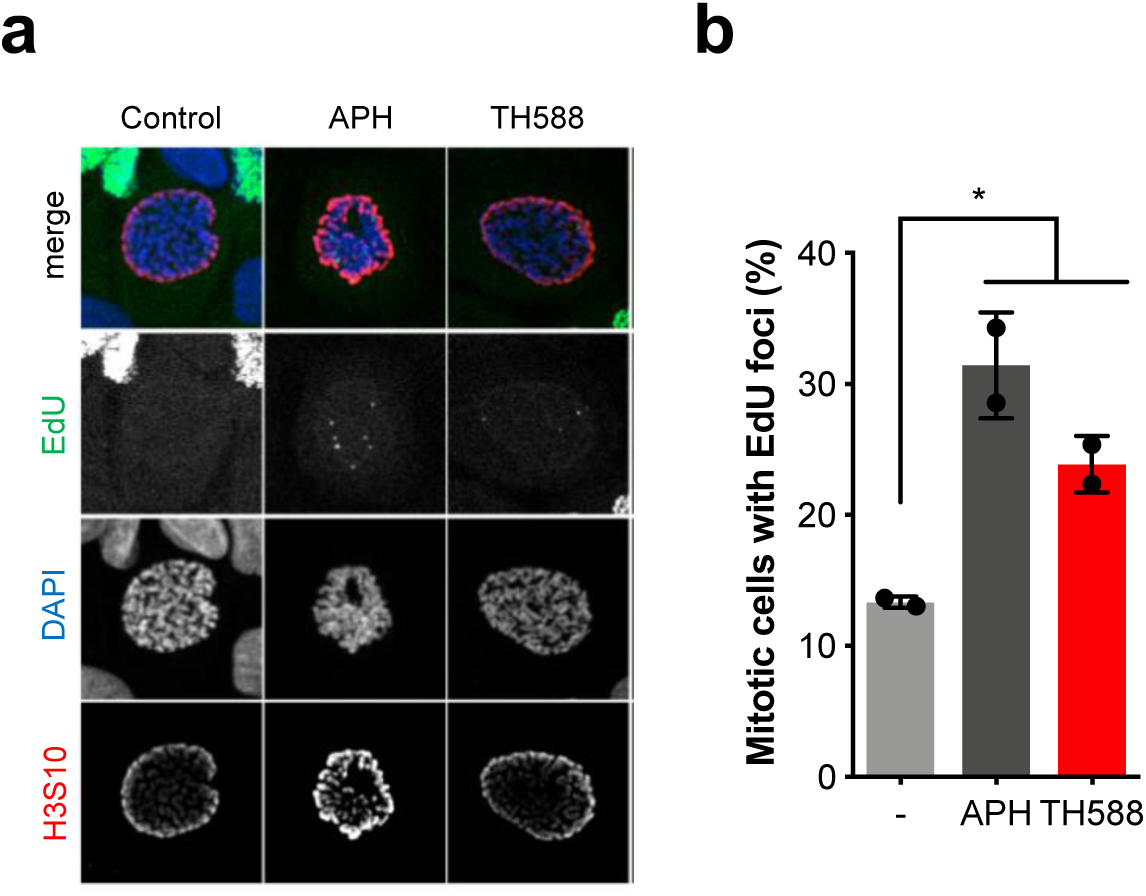
TH588 triggers mitotic replication. **a, b)** U2OS cells were treated with Cdk1 inhibitor RO3306 together with 0.4 μM aphidicolin (APH) or 5 μM TH588 for 16 h. Compounds were washed away and cells were released for 30 min in the presence of 10 μM EdU. EdU was labelled with Alexa-488-azide, mitotic cells with an H3-pS10 antibody, and DNA with DAPI. Representative images shown in (a), quantification of cells containing mitotic EdU foci shown in (b). Data shown as mean of >160 cells analysed per condition from 2 independent experiments, error bars indicate SD. Statistical test performed using Student’s t-test (* p<0.05).

### Spindle assembly checkpoint-mediated mitotic arrest is required for 8-oxodG accumulation and toxicity induced by TH588

Since replication in S phase is not greatly perturbed by TH588 treatment and instead this agent triggers mitotic DNA replication, we hypothesised that incorporation of 8-oxodGTP caused by TH588 into DNA may be dependent upon mitotic arrest, in order to generate ROS and provide time for mitotic DNA synthesis. To test this hypothesis we used an inhibitor to the mitotic kinase mps1 (reversine) that prevents the spindle assembly checkpoint (SAC) activation (30) in combination with TH588 treatment in U2OS cells. TH588 treatment induces a mitotic arrest (Figure 5a), and here we show that this arrest is dependent upon a functional SAC as reversin co-treatment ablates this mitotic arrest (Figure 5a). In accordance with our hypothesis, reversin treatment also prevents accumulation of TH588-induced genomic 8-oxodG and cell death (Figure 5b, c). Thus, this data demonstrates that SAC-dependent mitotic arrest is required for accumulation of genomic 8-oxodG and cell death following treatment with TH588.

**Figure 5.**
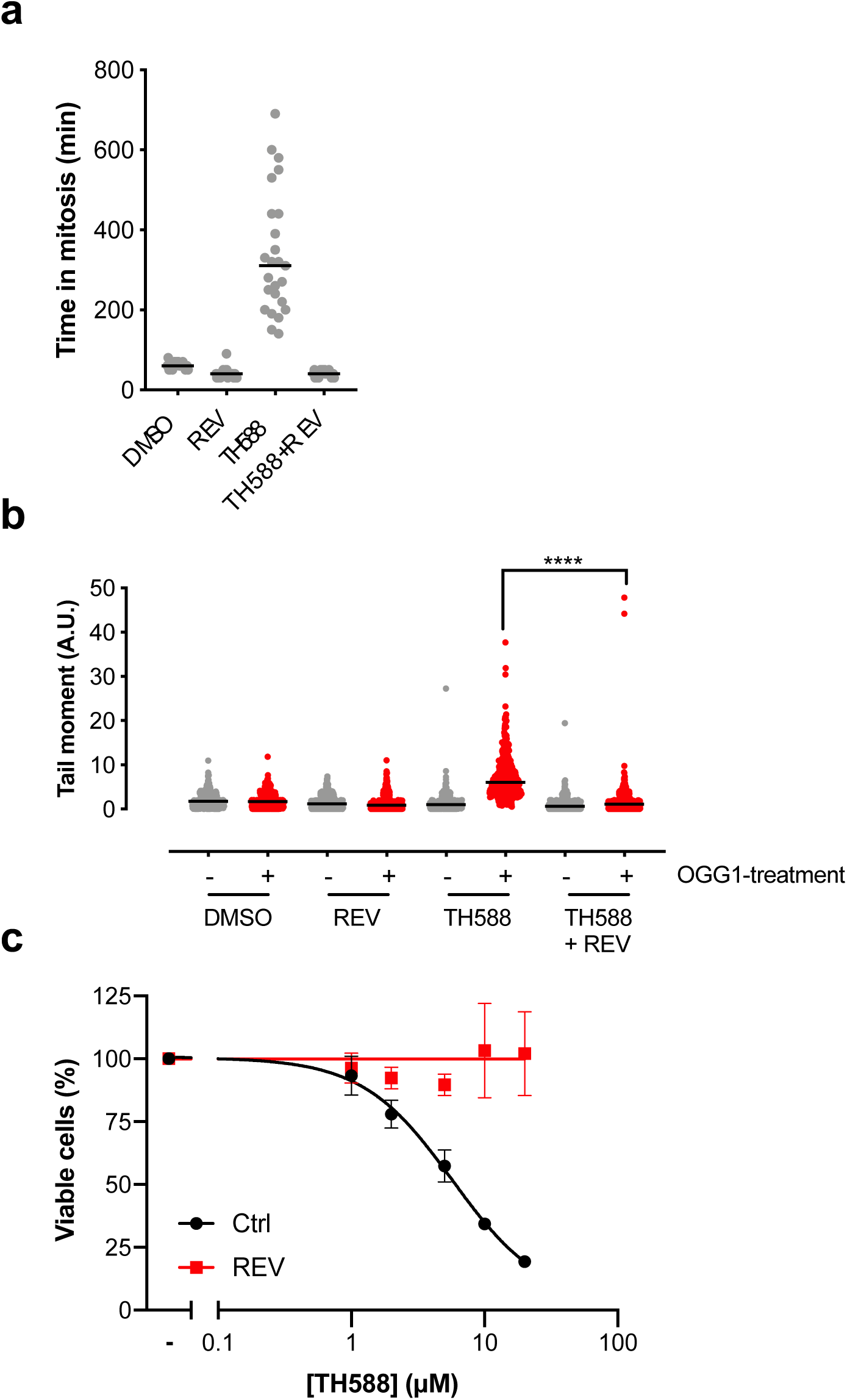
Mitotic arrest required for accumulation of genomic 8-oxodG following treatment with TH588. **a)** Quantification of the time in mitosis from the onset of pro-phase until the end of cytokinesis in U2OS-H2B-GFP cells treated with 10 μM TH588 and 0.5 μM Reversine for 24 h. Individual cells shown from a representative experiment, horizontal line indicates the mean. **b)** Relative tail moment in the modified comet assay of U2OS cells treated with 10 μM TH588 and 0.5 μM Reversine for 24 h. Data from two independent experiments shown with 400 cells analysed in total, horizontal line indicates the median, statistical testing was performed using the Mann-Whitney test (****, p<0.001). **c)** Viability assay of U2OS cells treated with TH588 with or without 0.5 μM Reversine for 72 h. Mean of 3 independent experiments shown, error bars indicate SD.

### Delayed mitotic progression in Pol δ EP cells treated with TH588, but not by anti-tubulin poisons

We hypothesised that additional mis-incorporation of oxidised nucleotides prior to, or during, mitotic entry could contribute to the mitosis-dependent cell death mechanism we observed. To test this we generated H2B-GFP expressing Pol δ replacement cells, and following TH588 treatment, monitored mitotic progression using live cell microscopy. In doxycycline-induced Pol δ WT cells, low-doses of TH588 induced a mitotic arrest, but the majority of cells could proceed into anaphase without apparent problems (Figure 6a, b; **Movie S1**). In contrast, the Pol δ EP expressing cells had a prolonged mitotic arrest in the presence of TH588 and underwent mitotic catastrophe, either becoming polynucleated or undergoing mitotic cell death (Figure 6a, b; **Movie S2**). These data suggest that oxidised dNTPs incorporated during mitotic replication can result in mitotic catastrophe.

**Figure 6.**
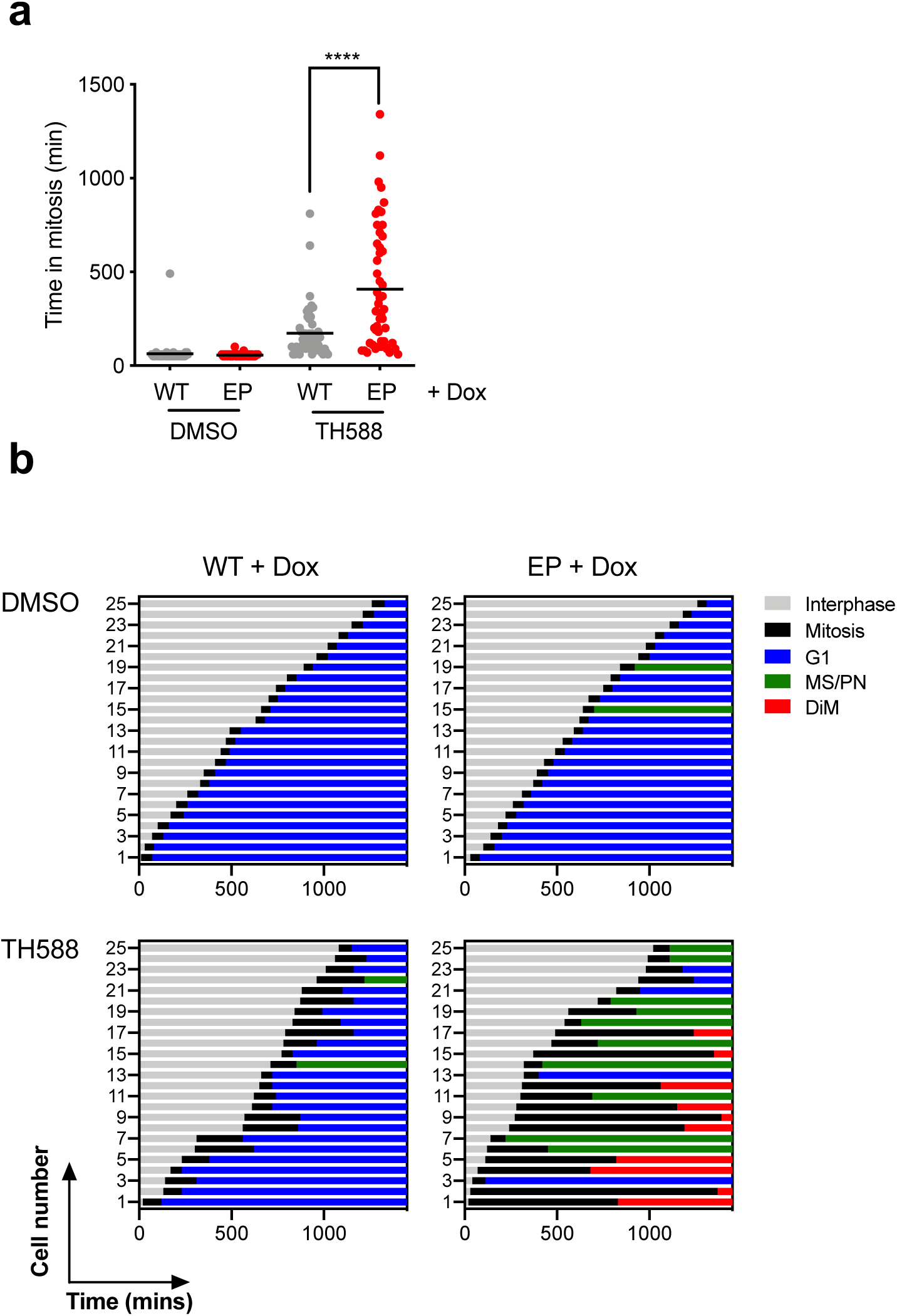
Expression of an error-prone variant of Pol δ results in a prolonged mitotic arrest and elevated mitotic cell death following TH588 treatment. **a)** Quantification of the time in mitosis from onset of pro-phase until end of cytokinesis. Data from two experiments, 50 cells analysed in total, horizontal line indicates the mean. Statistical testing performed with unpaired t-test (****p>0.001). **b)** Pol δ replacement-H2B-GFP cells (WT, wild type; EP, error-prone) were treated with 10 ng/ml doxycyclin for 4 days before treatment with 2.5 μM TH588 and subsequent initiation of the time-lapse microscopy after 30 min. Images were acquired every 10 min for 24 h. Individual cells were followed manually and scored for defects in mitosis including mitotic slippage/polynucleation (MS/PN), micronuclei formation (G1/MN), mitotic slippage (MS) or cell death during mitosis (DiM). Representative of two independent experiments shown.

To offer an alternative explanation, the mitotic defects induced by TH588 in Pol δ EP but not in WT expressing cells could potentially be a consequence of direct induction of ROS by the compounds, and Pol δ EP cells being able to incorporate 8-oxodGTP into DNA. Hence the effect may be independent of MTH1 inhibition by TH588. To test this, we triggered mitotic arrest and ROS using the tubulin poison paclitaxel and followed mitosis using live cell imaging. Interestingly, we found no difference in mitotic progression between Pol δ EP and WT expressing cells (Figure 7a, b), showing that a mitotic delay by itself without concomitant inhibition of MTH1 is insufficient to cause a differential response in Pol δ EP and WT expressing cells. To further support that MTH1 inhibition is important for the observed effect, Pol δ EP and WT cells were treated with other mitosis-targeted compounds, vincristine or an inhibitor to CENP-E, which resulted in no difference in mitotic progression between the two cell types (Figure 7c). Taken together, these data indicate that cancer cell death caused by TH588 is a combinatorial effect due to perturbation of microtubule dynamics and incorporation of oxidised nucleotides into the genome. Furthermore, mis-incorporation of oxidised dNTPs may alter mitotic progression.

**Figure 7.**
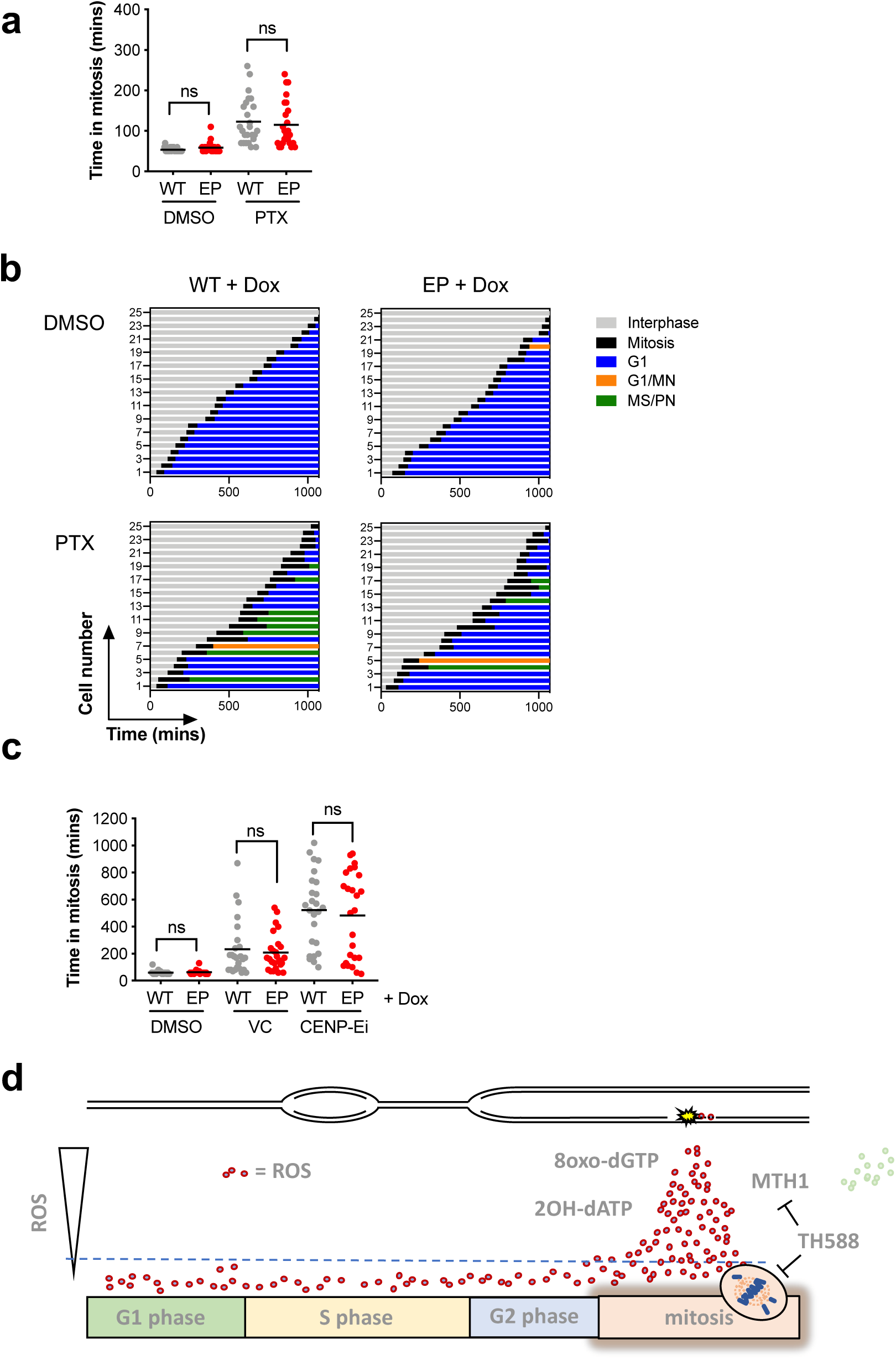
Expression of an error-prone variant of Pol δ does not result in prolonged mitotic arrest and elevated mitotic cell death induced by mitotic poisons. **a)** Quantification of the time in mitosis from the onset of pro-phase until the end of cytokinesis followed treatment with 2 nM Paclitaxel (PTX). Individual cells shown from a representative experiment, horizontal line indicates the mean. Statistical testing performed using unpaired t-test (ns, not significant). **b)** Time-lapse imaging of Pol δ replacement-H2B-GFP cells induced with 10 ng/ml doxycycline for 4 days prior to treatment with 2 nM Paclitaxel (PTX). Images were acquired every 10 min for 24 h. Individual cells were followed manually and scored for defects in mitosis including mitotic slippage/polynucleation (MS/PN), micronuclei formation (G1/MN), mitotic slippage (MS) or cell death during mitosis (DiM). Representative experiment shown. **c)** Quantification of the time in mitosis of Pol δ replacement-H2B-GFP cells induced with 10 ng/ml doxycycline for 4 days prior to treatment with 1 nM Vincristine or 20 nM CENP-Ei. Individual cells shown from a representative experiment, horizontal line indicates the mean. Statistical testing performed using unpaired t-test (ns, not significant). **d)** Mechanism of action of TH588. We propose MTH1 inhibition during S phase does not cause replication stress is related to overall low ROS levels also in cancer cells. Prolonged mitotic arrest causes mitophagy and ROS (28), likely above physiological levels (dotted line). We propose TH588 is causing ROS because of mitotic arrest mediated by directly interfering with tubulin polymerization or breaking MTH1-tubulin interactions. Through MTH1 inhibition, TH588 prevents 8-oxodGTP sanitization resulting in 8-oxodG incorporation into DNA during mitotic replication. We speculate cancer-specific toxicity of TH588 is related to replication stress associated in cancer, likely triggering mitotic replication, i.e., repair synthesis of under replicated regions.

## DISCUSSION

Here, we aimed to explore if and how oxidised dNTPs, incorporated into DNA, are toxic to cells. we used a previously established doxycycline-inducible replacement of endogenous Pol δ with an error-prone (L606G) variant (15), which we confirm did not greatly alter cell cycle progression or overall viability of cells. We demonstrate increased 8-oxodGTP incorporation in these cells, providing an experimental system by which to study incorporation of oxidative nucleotides into DNA and the resultant cellular consequences.

Since Pol δ is a main DNA polymerase one would expect a huge increase in 8-oxodG DNA levels and detrimental effects on replication in Pol δ EP cells exposed to MTH1i TH588, which we show is not the case. Since we observe minor alterations, and no generation of post-replication gaps, we suggest that the overall level of 8-oxodGTP is very low in the S phase of the cell cycle making MTH1 function redundant during this phase in unstressed conditions.

ROS is known to accumulate throughout the cell cycle, peaking in mitosis, and accordingly is induced following extended mitotic arrest (27,28). TH588 causes arrest in mitosis owing to a direct effect on tubulin (10) and likely also through disrupting the binding between MTH1 and tubulin in cells, resulting in reduced microtubule polymerisation rates (Gad et al., submitted). The discussion of the role of MTH1 on tubulin and in mitosis and the role of TH588 binding to tubulin is detailed elsewhere (Gad et al., submitted) and not the topic of this report. Nonetheless, TH588-induced arrest in mitosis is associated with an increase in ROS likely resulting in increased 8-oxodGTP levels in mitosis. Here, we report an increase in mitotic DNA replication following treatment with TH588 and also that the incorporation of 8-oxodGTP into DNA is dependent on an extended mitotic arrest, as pharmacologically reducing the time in mitosis reduced the incorporation of 8-oxodGTP as well as toxicity of TH588 (Figure 5).

A highly surprising finding is that Pol δ EP cells show a more pronounced delay in mitosis as compared to Pol δ WT cells suggesting incorporation of 8-oxodGTP into DNA is increasing the mitotic defect (Figure 6). Understanding how this work mechanistically will be challenging and should be addressed in future studies. An alternative explanation is that Pol δ EP cells would be prone to generate severe mitotic arrest owing to MTH1 and 8-oxodGTP independent effects. We argue this is not the case, as this mitotic arrest is not a phenomena observed generally following treatments with microtubule poisons (Figure 7).

Here, we propose a suitable model on the mechanism of action of TH588 (Figure 7d). TH588 does not lead to substantial obstruction of replication forks in S-phase due to low levels of ROS, which is supported by a recent study demonstrating elevated ROS levels in cancer cells primarily during G2 and particularly mitosis (27), which may also explain why other MTH1i that do not arrest cells in mitosis are non-toxic. ROS are accumulating upon mitotic arrest by TH588, mediated through a direct effect on tubulin and potentially also by disrupting MTH1 mitotic function and preventing the tubulin-MTH1 interaction (Gad et al., submitted). The nucleotide pool is oxidised in the presence of ROS in mitosis generating 8-oxodGTP, which is a substrate for MTH1 which prevents mitotic DNA damage. Upon inhibition of MTH1 8-oxodGTPase activity by TH588, 8-oxodGTP is incorporated into DNA during mitotic replication in cancer cells and enhances the mitotic arrest by a mechanism yet to establish.

## Supporting information

Move S2

## SUPPLEMENTARY DATA

Supplementary Data consists of one figure and two movies.

## ACKNOWLEDGEMENT

We thank Prof. Jircny (University of Zurich) for sharing Pol δ-replacement cell lines.

## AUTHOR CONTRIBUTIONS

T.H. and U.W.B. conceived and supervised the project. S.G.R., H.G., O.M., N.A., A.H., P.G., C.E.S. and T.H. designed, conducted and analysed experiments. S.G.R. and T.H. wrote the manuscript. All authors discussed results and approved the manuscript.

